# Angiotensin II Promotes Mitral Valve and Left Atrial Pathology in Fibrillin-1^C1041G/+^ Mice

**DOI:** 10.64898/2026.10.02.755732

**Authors:** Michael K. Franklin, Deborah A. Howatt, Alex C. Pettey, Jessica J. Moorleghen, Bryana M. Levitan, Adam E. Mullick, Jeffrey E. Saffitz, Hong S. Lu, Alan Daugherty

## Abstract

**Background:** Activation of the renin-angiotensin system (RAS) contributes to aortic diseases caused by fibrillin-1 haploinsufficiency (*Fbn1*^C1041G/+^). However, whether RAS activation influences mitral valve disease associated with FBN1 variants remains undefined. This study determined whether inhibition or augmentation of the RAS affects mitral valve pathology in *Fbn1*^C1041G/+^ mice.

**Methods:** Angiotensinogen (AGT) antisense oligonucleotides (ASOs) were administered for 12 weeks to inhibit the RAS in male and female *Fbn1*^C1041G/+^ mice. To augment the RAS, mice were infused with angiotensin II (AngII; 1,000 ng/kg/min) for 28 days. Mitral valve and heart function were monitored by ultrasound. At termination, hearts were dissected free and tissue sectioned to assess mitral valve leaflet thickness and pathology.

**Results:** Mitral valve leaflet thickness was significantly increased in *Fbn1*^C1041G/+^ mice compared to *Fbn1*^+/+^ littermates of both sexes. ASO-mediated AGT inhibition did not alter this phenotype despite markedly reduced plasma AGT concentrations. AngII infusion increased mitral valve leaflet thickness in *Fbn1*^C1041G/+^ mice with increased proteoglycan and collagen content, while exerting minimal effects in *Fbn1*^+/+^ mice. Twenty-five percent of AngII-infused *Fbn1*^C1041G/+^ mice died of non-vascular causes. Ultrasound showed mitral valve dysfunction in AngII-infused *Fbn1*^C1041G/+^ mice. Grossly expanded left atria were common in AngII-infused *Fbn1*^C1041G/+^ mice. These markedly enlarged left atria contained increased fibrosis, macrophage accumulation, and platelet-rich masses.

**Conclusions:** Mitral valves of *Fbn1*^C1041G/+^ mice did not respond to AGT inhibition but were markedly more susceptible to AngII infusion, with greater increases in thickness and compositional changes than mitral valves of *Fbn1*^+/+^ mice. In addition, AngII promoted mitral valve dysfunction that were associated with pronounced left atrial pathology.

**What are the Clinical Implications?:** Manipulation of fibrillin-1 in mice leads to vascular pathology that is attenuated by sustained inhibition of the renin-angiotensin system and exacerbated by its activation. However, whether modulation of the RAS influences mitral valve disease in mice harboring fibrillin-1 variants remains unknown. This study demonstrated that the presence of a missense variant of fibrillin-1 greatly augments the effects of renin-angiotensin system activation on mitral valve pathology compared with wild-type mice, with increased leaflet thickness and enhanced accumulation of proteoglycan and collagen. These structural changes are accompanied by mitral valve dysfunction and pronounced left atrial pathology, likely attributable to angiotensin II-induced mitral valve prolapse. These findings provide an experimental basis for determining whether inhibition of the renin-angiotensin system attenuates the development or progression of mitral valve dysfunction in humans with fibrillin-1 variants.

## INTRODUCTION

Fibrillin-1 (FBN1) variants in individuals with Marfan syndrome are commonly studied in the context of proximal thoracic aortic aneurysms. Consequently, experimental studies in mice have focused largely on interventions that attenuate disease in this aortic region. Inhibition of the renin-angiotensin system reduces thoracic aortic disease in mice with FBN1 manipulation, as demonstrated by suppression of angiotensinogen (AGT) synthesis with antisense oligonucleotides (ASOs)^1^ and by pharmacological or genetic inhibition of angiotensin II (AngII) signaling through AT1a receptors in *Fbn1*^C1041G/+^ mice.^1–4^ Conversely, activation of the renin-angiotensin system through chronic subcutaneous AngII infusion augments vascular pathology in mice with genetic manipulation of FBN1.^5, 6^

In addition to aortopathy, *FBN1* variants are frequently associated with mitral valve defects in humans.^7–9^ Consistent with these clinical manifestations, *Fbn1*^C1041G/+^ mice develop mitral valve pathology.^10–13^ However, whether the renin-angiotensin system contributes to mitral valve disease in this model remains undefined. Defining this relationship in *Fbn1*^C1041G/+^ mice may provide translational insight into mechanisms that drive mitral valve disease in Marfan syndrome.

The renin-angiotensin system can be manipulated by blocking AT1 receptors, inhibiting renin or angiotensin-converting enzyme activity, suppressing AGT synthesis, or augmenting the pathway with AngII infusion. ASO-mediated AGT suppression provides sustained inhibition because of the protracted half-life of ASOs and the central role of AGT as the precursor of all angiotensin peptides.^1, 14^ In contrast, subcutaneous AngII infusion is widely used to define the contribution of enhanced AngII signaling to cardiovascular diseases, including hypertension,^15^ cardiac dysfunction,^16^ and aortopathy.^17^ Together, these complementary approaches provide a strategy to determine whether reduced or enhanced activity of the renin-angiotensin system modifies mitral valve pathology in *Fbn1*^C1041G/+^ mice. Building on this rationale, this study determined whether sustained AGT suppression attenuates myxomatous mitral valve pathology and whether AngII infusion exacerbates mitral valve pathology in *Fbn1*^C1041G/+^ mice compared to *Fbn1*^+/+^ littermates.

## METHODS

### Data Availability

Detailed materials and methods are available in this manuscript and the supplemental material. Numerical data are presented in the Supplemental Data File. Studies were designed and performed in accordance with recently published recommendations.^18^

### Mice

Breeding pairs of *Fbn1*^+/+^ and *Fbn1*^C1041G/+^ mice (stock #012885) were obtained from The Jackson Laboratory. Male *Fbn1*^C1041G/+^ mice were bred with female *Fbn1*^+/+^ mice to generate *Fbn1*^+/+^ and *Fbn1*^C1041G/+^ littermates. After weaning, littermates were separated by sex and randomly assigned to study groups. Mice were housed with up to 5 animals per cage and maintained on a 14:10-hour light:dark cycle. They were fed Teklad Irradiated Global 18% Protein Rodent Diet (Diet # 2918, Inotiv) *ad libitum* and had *ad libitum* access to water through a Lixit system. Bedding was provided by P.J. Murphy (Coarse Sani-Chips) and changed biweekly throughout the study. Room temperature was maintained at 21 ± 2 °C, and room humidity was maintained at approximately 50%.

Mice were monitored daily for health, weighed weekly, and underwent necropsy to determine the cause of death if any study mice died prematurely before study termination. All protocols were approved by the University of Kentucky IACUC (Protocol # 2018-2967).

### Genotyping

*Fbn1*^C1041G/+^ was genotyped using forward primer (5’-CTCATCATTTTTGGCCA GTTG-3’) and reverse primer (5’-GCACTTGATGCACATTCACA-3’) based on the protocol described by The Jackson Laboratory.^1^ *Fbn1*^+/+^ generated a 164 bp product. *Fbn1*^C1041G/+^ generated a 212 bp (mutant) product and a 164 bp (wild type) product. Post-termination validation genotyping was performed by Transnetyx in a blinded manner.

### Administration of Antisense Oligonucleotides or Angiotensin II

Angiotensinogen antisense oligonucleotides (AGT ASO #1095847-9; 5’-ATCATTTATTCTCGGT-3’) were provided by Ionis Pharmaceuticals. AGT ASO was administered once weekly by subcutaneous injection for 12 weeks, beginning at 4-5 weeks of age, at a dose of 40 mg/kg/week, as described previously.^1, 19^ AGT suppression was confirmed by measuring plasma concentrations using a mouse AGT ELISA (Catalog #: 27103; IBL).^19^

In a separate study designed to determine whether AngII augments mitral valve pathology, AngII was infused in mice 8 - 10 weeks of age at 1,000 ng/kg/min using osmotic pumps (Alzet model 1004) implanted subcutaneously.^20, 21^

### Necropsy

Necropsies were performed immediately after carcasses were found to determine the cause of death. Vascular rupture was defined as the presence of extravascular blood accumulated within either the thoracic or abdominal cavity. Non-vascular-related death was defined by the absence of extravascular blood in either cavity. Cardiac tissues were collected at study termination from all mice that survived to the endpoint.

### Pathology

For histological analysis, hearts, including the left atrial appendage, were dissected free and immersed in neutral buffered formalin (10% wt/vol) overnight. Tissues were then dehydrated through a graded series of ethanol concentrations and embedded in paraffin wax. Tissue sections (5 μm) were collected using a microtome. Paraffin-embedded sections were deparaffinized using limonene (Cat # 6533A, Medical Chemical Corporation).

Histological staining, including hematoxylin and eosin, Masson’s trichrome, picrosirius red and fast green, and alcian blue and nuclear red was performed to visualize mitral valve and atrial pathologies and collagen fibers. For immunostaining, deparaffinized sections were incubated with H2O2 for 10 minutes at room temperature, followed by antigen retrieval for 20 minutes at 98 °C. Nonspecific binding sites were blocked using goat serum for 20 minutes at room temperature. Sections were incubated with primary antibodies overnight at 4 °C. Goat anti-rabbit IgG conjugated with horseradish peroxidase was used as the secondary antibody, and NovaRED was used as the chromogen. Reagent information is listed in Major Resources Tables II–IV. Histological and immunostained tissue sections were imaged using an Axioscan 7 (Zeiss) or Nikon Eclipse Ni microscope and analyzed using ZEN v3.1 blue edition (Zeiss) or NIS-Elements AR 5.11.03 software (Nikon Instruments Inc.).

### Measurement of Mitral Valve Thickness

To visualize the mitral valve leaflets, each whole heart was oriented along the anatomical long axis. Longitudinal serial sections were collected from the anterior to posterior aspects through the center of the mitral valve and left ventricular outflow tract. Hearts were sectioned carefully to preserve the mitral annulus, anterior and posterior mitral leaflets, chordae tendineae, and papillary muscles. The sectioning plane was maintained consistently across all specimens to minimize variability in morphometric measurements.

Sections containing the maximal cross-sectional area of the mitral valve leaflets were selected for analysis and stained with H&E. Anterior and posterior leaflet thicknesses were measured at the maximal midpoint of each leaflet using a line drawn perpendicular to the leaflet long axis. For each heart, mitral valve thickness was measured in 4-8 sections spaced approximately 150 μm apart across the central portion of the mitral valve, and the maximal value was used for statistical analysis. All measurements were verified by an investigator blinded to experimental groups using NIS-Elements AR 5.11.03 software (Nikon Instruments Inc.).

### Ultrasound

Cardiac structure and function were assessed by echocardiography using a Vevo 3100 ultrasound system equipped with an MS550 transducer (FUJIFILM VisualSonics Inc.). Mice were placed on a heated platform maintained at 37 °C to prevent hypothermia and anesthetized with isoflurane (1-2% vol/vol) to maintain heart rates between 400 and 550 beats/min.

To determine cardiac function, standard two-dimensional B-mode images were acquired from parasternal long-axis and short-axis views. Three consecutive M-mode recordings were obtained from the short axis bisecting the ventricle at the level of the papillary muscles to measure left ventricular internal diameter in diastole (LVIDd), left ventricular posterior and anterior wall thicknesses (LVPWd and LVAWd), and left ventricular mass. Left ventricular ejection fraction and fractional shortening were calculated using the manufacturer’s analysis software. Mitral valve function was assessed using parasternal long-axis and apical four-chamber views when feasible. Color Doppler imaging was performed to evaluate mitral regurgitation when indicated.

Images were analyzed offline using Vevo LAB software (FUJIFILM VisualSonics) by an investigator blinded to genotype and treatment group. Measurements from at least three consecutive cardiac cycles were calculated for each animal and used as the mean value for statistical analysis.

### Statistical Analyses

Statistical analyses were performed using SigmaPlot version 16. Data are presented as mean ± SEM for parametric analyses and as median with interquartile range (25th and 75th percentiles) for nonparametric analyses. For raw or log-transformed data with n>5/group, normality and homogeneity of variance were assessed using the Shapiro-Wilk and Brown-Forsythe tests, respectively. For comparisons between two groups that met both assumptions, Student’s *t* test was used. For data that passed the normality test but failed the equal variance test, Welch’s t test was used. For comparisons among multiple groups that met the assumptions of both normality and equal variance, two-way analysis of variance was performed, followed by Holm-Sidak post hoc tests. Kruskal-Wallis or rank-sum tests were used for nonparametric comparisons when data failed either the normality or equal variance test, followed by Dunn’s method for post hoc testing when appropriate. P<0.05 was considered statistically significant. Data are presented in either a parametric or non-parametric format consistent with the statistical analysis performed.

## RESULTS

### Inhibition of AGT synthesis did not attenuate fibrillin-1 haploinsufficiency-induced mitral valve leaflet thickening

Because mitral valve thickening in *Fbn1*^C1041G/+^ mice has been reported to become evident at 2 months of age,^11^ we initiated AGT ASO administration at 4-5 weeks of age, before overt valve thickening developed. AGT ASO reduced plasma AGT concentrations by more than 90% in both male and female mice after 12 weeks of administration (Figure S1). In vehicle-administered mice, plasma AGT concentrations were comparable between *Fbn1*^+/+^ and *Fbn1*^C1041G/+^ genotypes.

Heart tissue regions containing the mitral valves were sectioned longitudinally from the anterior aspect throughout the entire leaflets. These sections showed that mitral valve leaflets were thicker at study termination at 4 months of age in male and female *Fbn1*^C1041G/+^ mice than in their *Fbn1*^+/+^ littermates (Figure 1A and 1B). However, AGT ASO administration did not significantly affect leaflet thickness in either male or female *Fbn1*^C1041G/+^ mice compared with vehicle administration in mice of both sexes (Figure 1A and 1B).

**Figure 1.**
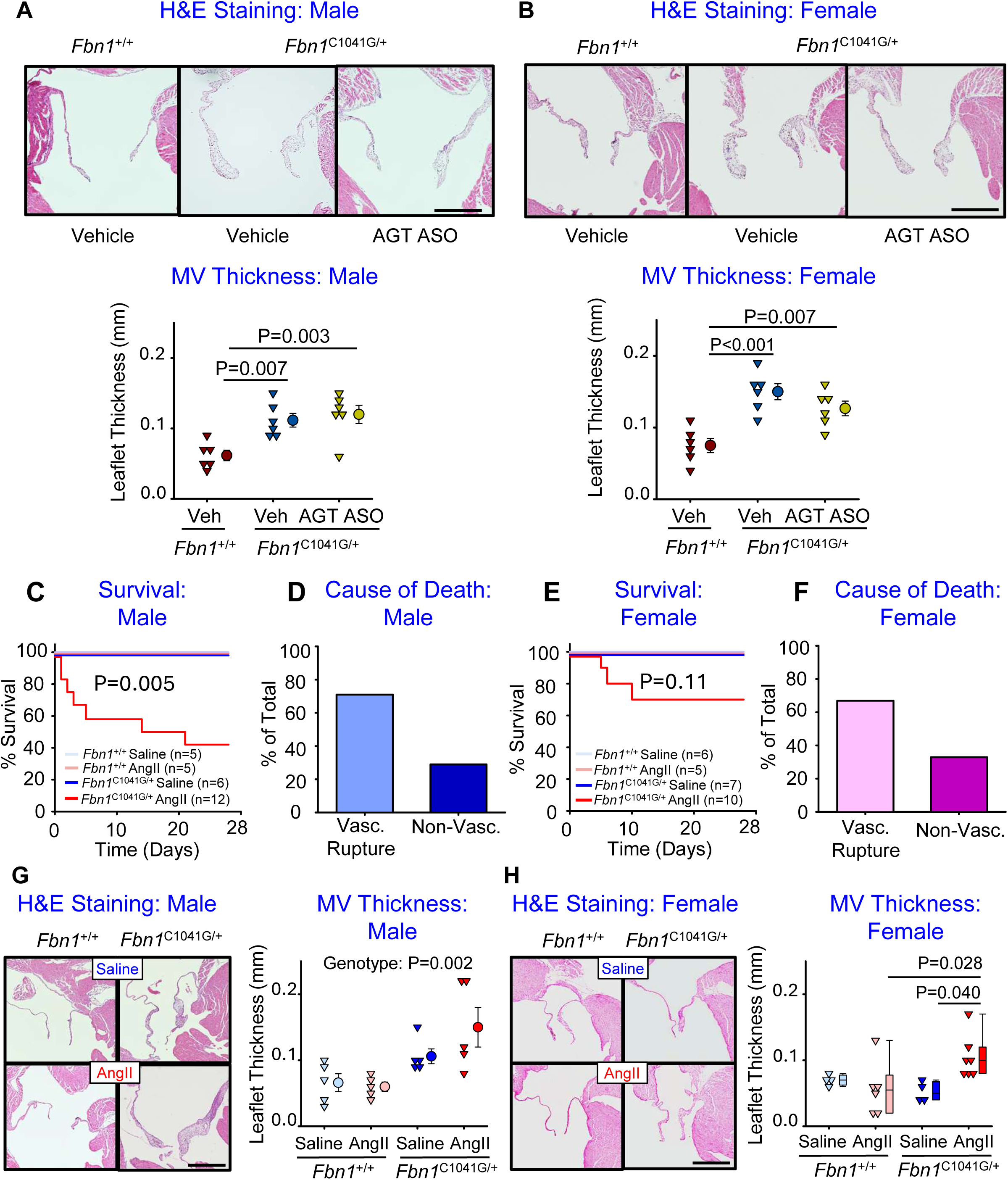
Mitral valve leaflet thickening in *Fbn1*^C1041G/+^ mice was unaffected by AGT inhibition while increased by AngII infusion. **A and B.** *Fbn1*^+/+^ mice were administered PBS (vehicle), and *Fbn1*^C1041G/+^ mice were administered vehicle or AGT ASO by subcutaneous injection once weekly for 12 weeks, beginning at 4-5 weeks of age. Mitral valve thickness was visualized and quantified using hematoxylin/eosin (H&E) stained heart sections. n=6/group. Data were analyzed using one-way ANOVA followed by the Holm-Sidak post hoc test. **C-H.** Eight-to 10-week-old *Fbn1*^+/+^ and *Fbn1*^C1041G/+^ mice were infused with either saline or AngII (1,000 ng/kg/min) for 4 weeks. P values in **C and E** were determined by pairwise log-rank analysis. **G and H.** Mitral valve thickness was visualized and quantified using hematoxylin and eosin-stained tissue sections. n=5/group. Data were analyzed using two-way ANOVA with the Holm-Sidak method. Veh, vehicle; Vasc, vascular; Non-Vasc, non-vascular. Scale bars equal 500 µm.

### AngII infusion increased mitral valve leaflet thickness in *Fbn1*^C1041G/+^ mice

To determine whether activation of the renin-angiotensin system exacerbated mitral valve pathology, AngII was infused at a rate of 1,000 ng/kg/min for 4 weeks. AngII infusion led to substantial mortality, with deaths occurring in 58% of male and 30% of female *Fbn1*^C1041G/+^ mice (Figure 1C and 1E). Consistent with our previous findings,^5^ most deaths were attributable to aortic rupture; however, more than 20% occurred in the absence of apparent aortic rupture in both male and female *Fbn1*^C1041G/+^ mice (Figure 1D and 1F, and Table S1). To assess the effects of AngII on mitral valve structure, we quantified leaflet thickness at the end of the study and performed multiple histological stains. AngII infusion had no effect on mitral valve thickness in male and female *Fbn1*^+/+^ mice compared to saline-infused mice (Figure 1G and 1H). In contrast, AngII infusion for 4 weeks starting at 8-10 weeks of age significantly increased leaflet thickness in female *Fbn1*^C1041G/+^ mice compared to saline-infused mice (Figure 1G and 1H). Staining with picrosirus red and Alcian blue illustrated increased collagen and proteoglycans, respectively, in mitral valve leaflets and annuli of *Fbn1*^C1041G/+^ mice infused with AngII (Figure S3 and S4), affecting both the anterior and posterior leaflets (Figure S5 and S6).

### AngII infusion induced mitral valve dysfunction and left atrial pathology in Fbn1^C1041G/+^ mice

As described above, AngII-infused *Fbn1*^C1041G/+^ mice that died before study termination, without evident aortic rupture, exhibited striking enlargement of the left atrium. Even among mice that survived to the endpoint, many left atria were grossly enlarged and engorged (Figure 2 and Figure S2), suggesting possible left atrial bleeding. Given these prominent gross abnormalities, we assessed the composition of left atrial tissue. In saline-infused *Fbn1*^+/+^ mice, the left atrium was small and translucent, located at the superior aspect of the left ventricle, and contained minimal collagen content with few discernible macrophages (Figure 2A). In *Fbn1*^C1041G/+^ mice that survived 28 days of AngII infusion, the left atria were frequently enlarged, with more evident fibrosis and macrophage accumulation (Figure 2B). In mice that died without apparent aortic rupture, left atrial enlargement was even more pronounced (Figure 2C). Fibrosis was modestly increased along the outer tissue margins and colocalized with macrophage accumulation. Notably, the most distinctive feature was a white mass occupying most of the left atrium that stained positive for CD41, indicating a predominance of platelet accumulation.

**Figure 2.**
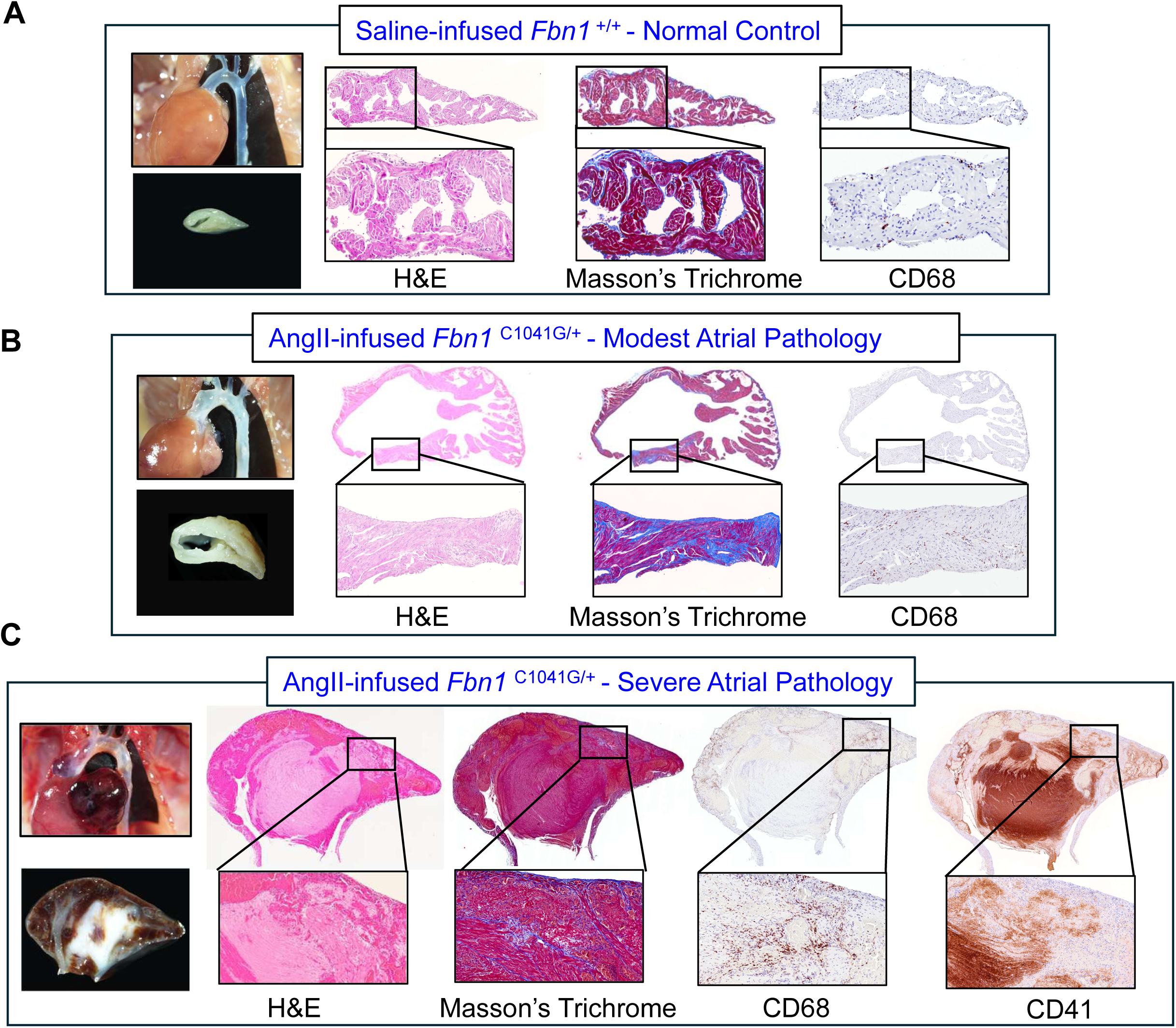
Chronic AngII infusion provoked mitral valve dysfunction and left atrial pathology in *Fbn1*^C1041G/+^ mice. Male *Fbn1*^+/+^ and *Fbn1*^C1041G/+^ mice were infused with saline or AngII (1,000 ng/kg/min) for 4 weeks. **A-C.** *In situ* images and gross cross-sections of left atria are shown with corresponding tissue sections stained with hematoxylin and eosin (H&E) or Masson’s trichrome, or immunostained for CD68 or CD41. **A.** Normal left atrium from a saline-infused *Fbn1*^+/+^ mouse. **B and C.** Pathological left atria from AngII-infused *Fbn1*^C1041G/+^ mice. Scale bars equal 100 µm.

Because of the striking pathological changes in left atria, we subsequently performed ultrasound imaging to determine whether we could observe any functional changes in the mitral valve. As noted in Video S1A, normal valve closing was detected in *Fbn1*+/+ mice. Both posterior and anterior mitral valve prolapse and regurgitation were detected in *Fbn1*^C1041G/+^ mice (Video S1B). The changes were more pronounced during AngII infusion into *Fbn1*^C1041G/+^ mice, with the presence of posterior and anterior mitral valve prolapse that was associated with enlarged left atrium and a flailing posterior mitral valve with an engorged left atrium (Video S1C & D).

AngII infusion increased left ventricular mass in male *Fbn1*^C1041G/+^ mice compared with their wild-type controls, whereas this parameter did not differ significantly in female mice. No other cardiac function differences were noted by ultrasound (Table S2).

## DISCUSSION

Mitral valve disease is a common manifestation of Marfan syndrome and is recapitulated in *Fbn1*^C1041G/+^ mice, which develop myxomatous mitral valve pathology.^10–13^ The present study used complementary approaches to determine whether reduced or enhanced activity of the renin-angiotensin system changed this phenotype. Sustained suppression of AGT synthesis markedly reduced plasma AGT concentrations but did not attenuate mitral valve leaflet thickening in either male or female *Fbn1*^C1041G/+^ mice. AngII infusion also had no effect on mitral valve thickness in *Fbn1*^+/+^ mice. Conversely, chronic AngII infusion produced increased mitral valve leaflet thickness in female *Fbn1*^C1041G/+^ mice and caused marked increases in proteoglycan and collagen content. AngII also provoked a striking left atrial phenotype characterized by marked left atrial enlargement and platelet-rich pathology. Together, these findings indicate that in *Fbn1*^C1041G/+^ mice, enhanced AngII signaling increases mitral valve pathology and enhances susceptibility to left atrial pathology.

Previous studies have implicated TGFβ-dependent mechanisms, inflammatory cell recruitment, and Wnt signaling in development and progression of myxomatous mitral valve disease in *Fbn1*^C1041G/+^ mice.^10–13^ In contrast, the contribution of the renin-angiotensin system to mitral valve pathology has been undefined. This study used AGT ASO administration, which provided a robust investigation of this pathway, because AGT is the precursor for all angiotensin peptides, and its suppression produces sustained inhibition of the renin-angiotensin system activity.^1, 19, 22^ Despite a profound effect of AGT suppression on thoracic aortopathy in *Fbn1*^C1041G/+^ mice,^1–4^ this approach had no discernible effect on mitral valve thickness in these mice. Several considerations may explain the lack of effect of AGT suppression on mitral valve pathology. First, leaflet thickening in the vehicle-administered *Fbn1*^C1041G/+^ mice was modest at the age examined, which may have limited the ability to detect attenuation. Second, mitral valve remodeling may occur during developmental or early postnatal windows that preceded initiation of ASO administration. Third, tissue-specific mechanisms within the valve that are independent of angiotensin II activation may be dominant contributors to this phenotype. Therefore, although our studies using systemic AGT suppression did not attenuate mitral valve disease in *Fbn1*^C1041G/+^ mice, further studies would benefit from initiating the drug at increasing ages of more advanced disease and having more protracted interval in ASO administration.

In contrast to AGT inhibition, AngII infusion enhanced mitral valve pathology. AngII infusion is widely used to induce or promote vascular and cardiac pathology, including aortic disease in susceptible mouse strains.^5, 23–25^ In the present study, AngII infusion did not alter mitral valve leaflet thickness in *Fbn1*^+/+^ mice. The high death rate in the male *Fbn1*^C1041G/+^ mice led to limited statistical power to determine the effects on valve thickness. However, AngII infusion increased collagen and proteoglycan deposition in mitral valves of *Fbn1*^C1041G/+^ mice. These data are consistent with the interpretation that enhanced AngII signaling alone in wild type mice was not sufficient to produce a robust mitral valve phenotype. The modest sex difference in mitral valve thickness should be interpreted cautiously because mortality during AngII infusion reduced the number of mice available for endpoint valve measurements, particularly among male *Fbn1*^C1041G/+^ mice.

The most striking finding of this study was the profound left atrial pathology induced by AngII infusion in *Fbn1*^C1041G/+^ mice. Mice that died without overt aortic rupture and mice that survived to termination frequently exhibited markedly enlarged left atria. Histological analyses showed fibrosis, macrophage accumulation, and large CD41-positive platelet-rich masses occupying much of the atrial lumen. A major unresolved issue is whether the modest structural changes in the mitral valve were sufficient to impair valve function and result in severe left atrial pathology. We attempted to assess mitral valve inflow and regurgitation by ultrasound during saline or AngII infusion, but these parameters could not be distinguished reliably in AngII-infused mice. However, direct visualization of the mitral valve demonstrated the presence of prolapse in the anterior and posterior leaflets. A flailing posterior mitral valve was also detected in AngII-infused *Fbn1*^C1041G/+^ mice associated with an engorged left atrium. Therefore, these data are consistent with AngII infusion into *Fbn1*^C1041G/+^ mice precipitating mitral valve dysfunction directly contributing to left atrial pathology. The left atrium has the highest abundance of FBN1 compared to other heart chambers, which may contribute to the location of the pathology.^26^ Alternatively, AngII may have acted directly on atrial tissue, leading to atrial remodeling independent of primary mitral valve dysfunction. This latter possibility is supported by emerging evidence in humans that cardiac manifestations beyond proximal aortic disease, including increased left atrial size without overt left ventricular dysfunction, occur in patients with Marfan syndrome.^27^

These findings have translational relevance because management of Marfan syndrome has traditionally focused on preventing proximal thoracic aortic expansion, whereas non-aortic cardiovascular manifestations are increasingly recognized.^7–9, 27–29^ Angiotensin receptor blockers and angiotensin-converting enzyme inhibitors are commonly used in patients with Marfan syndrome to limit aortic growth; however, their effects on mitral valve disease and atrial remodeling remain less well defined. The present findings demonstrate that systemic AGT suppression alone does not attenuate modest mitral valve thickening in *Fbn1*^C1041G/+^ mice, whereas increased AngII can provoke severe atrial pathology in this genetic background. Further studies focusing on pathological and mechanistic analyses of atrial tissues will be needed to determine whether this atrial pathology develops secondary to mitral valve dysfunction, results from direct AngII effects on FBN1-disrupted atrial myocardium or reflects a combination of both mechanisms.

## Non-standard Abbreviations

AGT: Angiotensinogen
AngII: Angiotensin II
ASO: Antisense oligonucleotides
FBN1: Fibrillin-1

## Acknowledgments

These studies were facilitated by the University of Kentucky Light Microscopy Core (RRID: SCR_ 026405).

## Supplemental Materials

### SUPPLEMENTAL FIGURE LEGENDS

**Figure S1.**
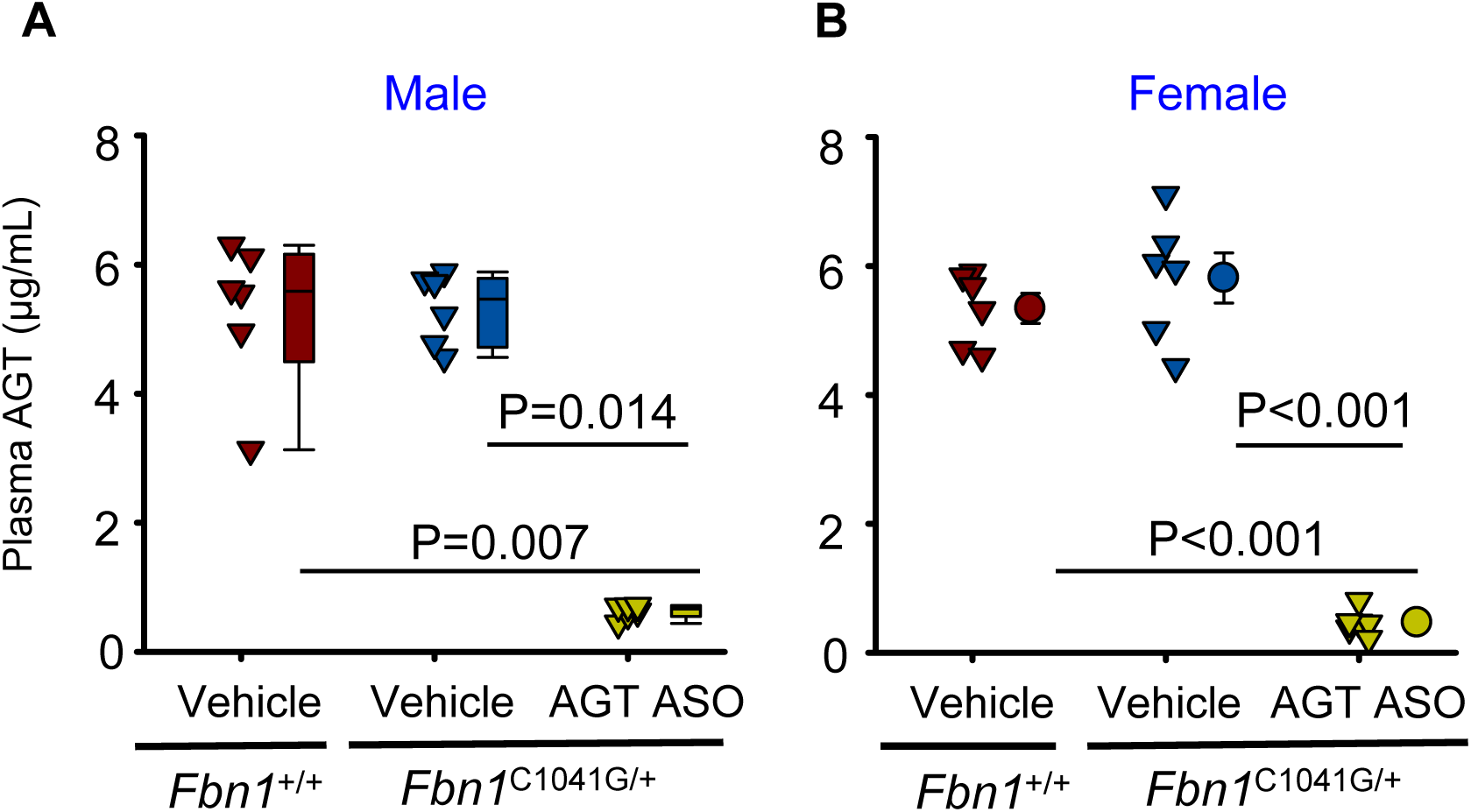
Inhibition of AGT synthesis suppressed plasma AGT concentrations. *Fbn1*^+/+^ mice were administered PBS (vehicle), and *Fbn1*^C1041G/+^ mice were administered vehicle or AGT ASO by subcutaneous injection once weekly for 12 weeks, beginning at 4 weeks of age. Plasma AGT concentrations were determined at the endpoint. Data were analyzed using Kruskal-Wallis one-way ANOVA on ranks followed by Dunn’s post hoc test for A and one-way ANOVA followed by Holm-Sidak post hoc test for B. n=6/group.

**Figure S2.**
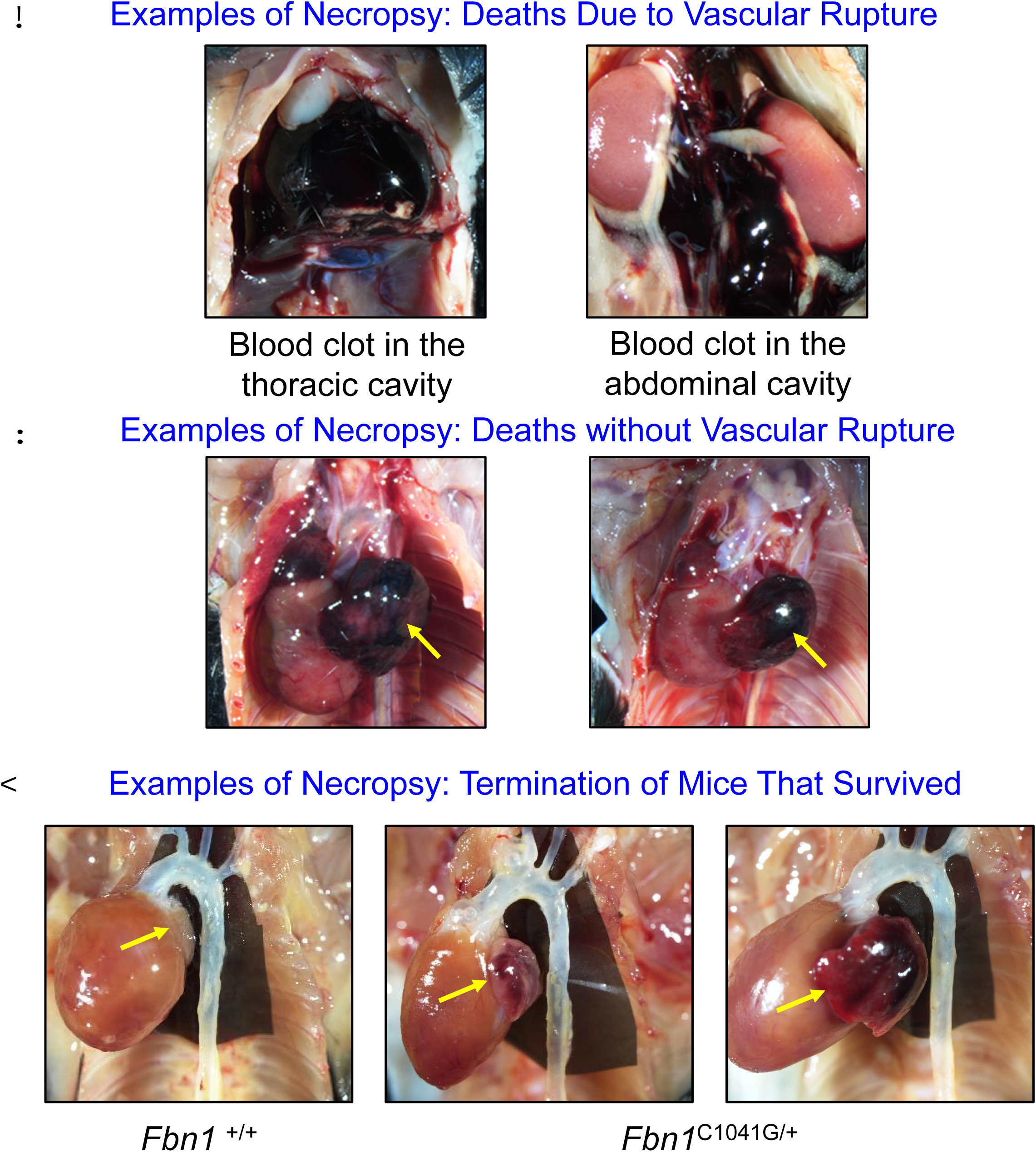
Chronic AngII infusion into *Fbn1*^C1041G/+^ mice provoked left atrial pathology. Male and female *Fbn1*^+/+^ and *Fbn1*^C1041G/+^ mice were infused with AngII (1,000 ng/kg/min) for 4 weeks. In situ images in A and B are from mice that died before the endpoint. In situ images in C show hearts, including left atria, from *Fbn1*^+/+^ and *Fbn1*^C1041G/+^ mice. Yellow arrows indicate the left atria.

**Figure S3.**
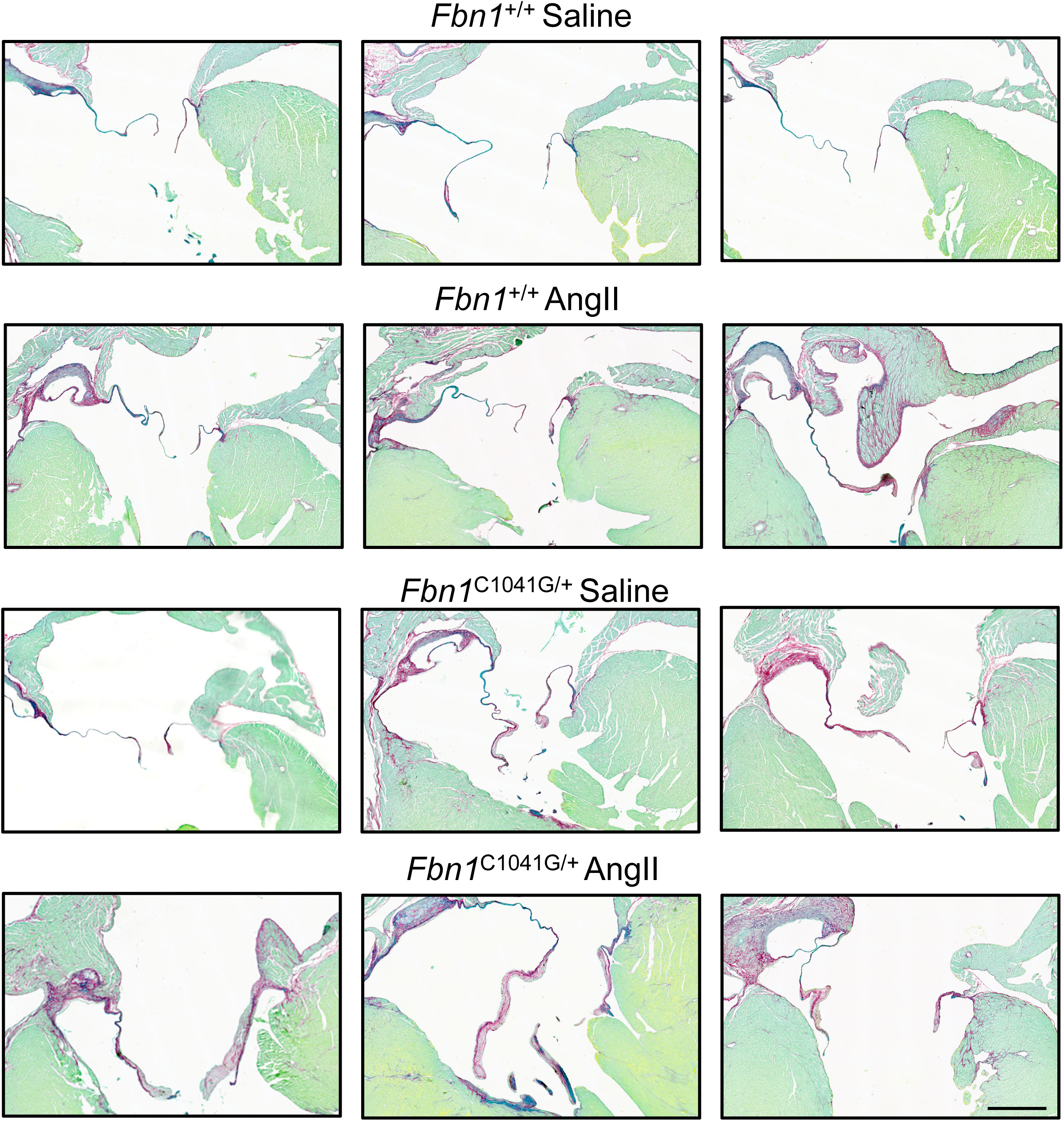
Chronic AngII infusion increased collagen deposition in mitral valves. Examples are for *Fbn1*^+/+^ and *Fbn1*^C1041G/+^ mice were infused with either saline or AngII (1,000 ng/kg/min) for 4 weeks. Tissue sections stained with picrosirius red and fast green. Scale bar equals 500 µm.

**Figure S4.**
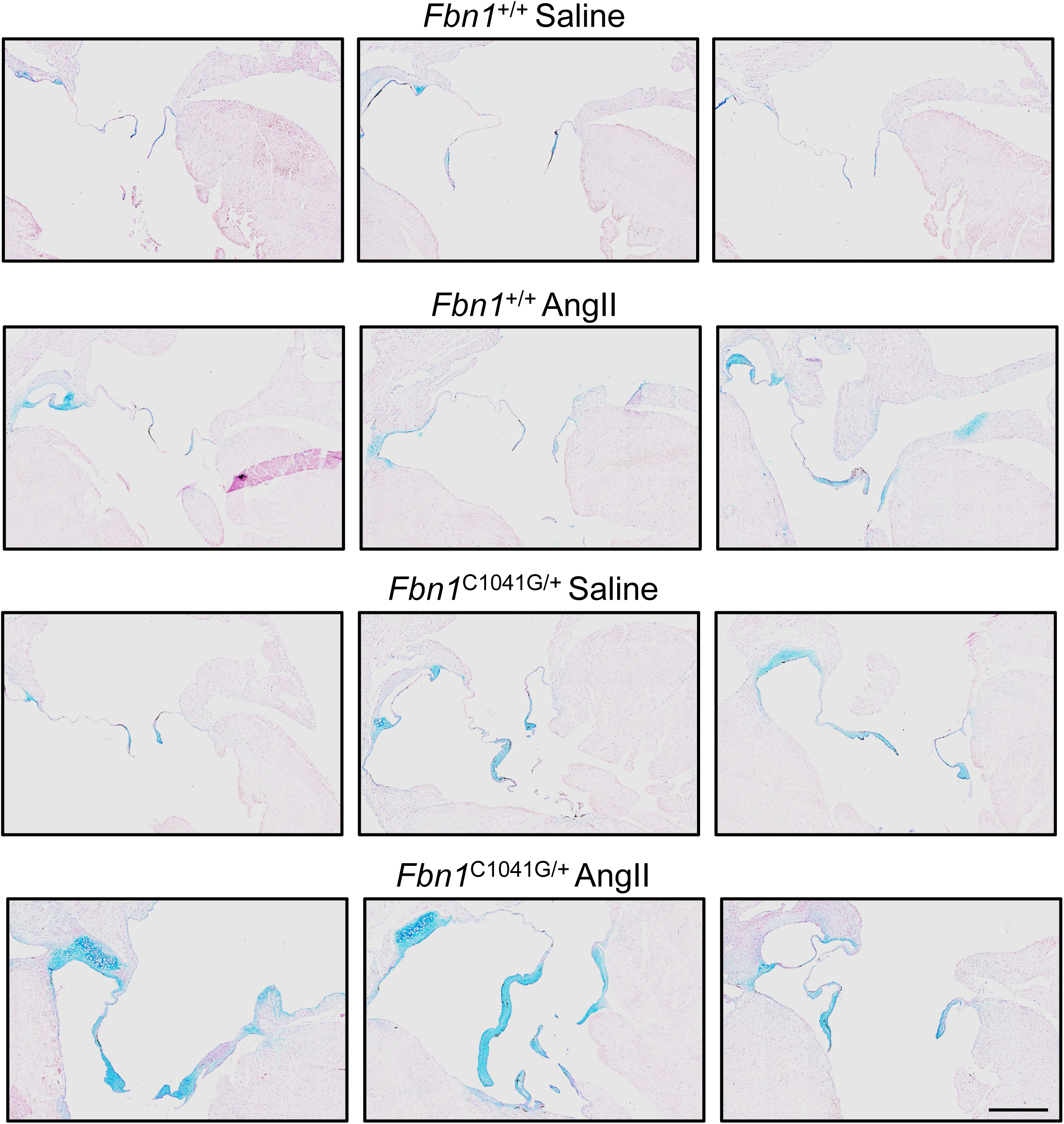
Chronic AngII infusion increased proteoglycan deposition in mitral valves. Examples are for *Fbn1*^+/+^ and *Fbn1*^C1041G/+^ mice were infused with either saline or AngII (1,000 ng/kg/min) for 4 weeks. Tissue sections stained with alcian blue and nuclear red. Scale bar equals 500 µm.

**Figure S5.**
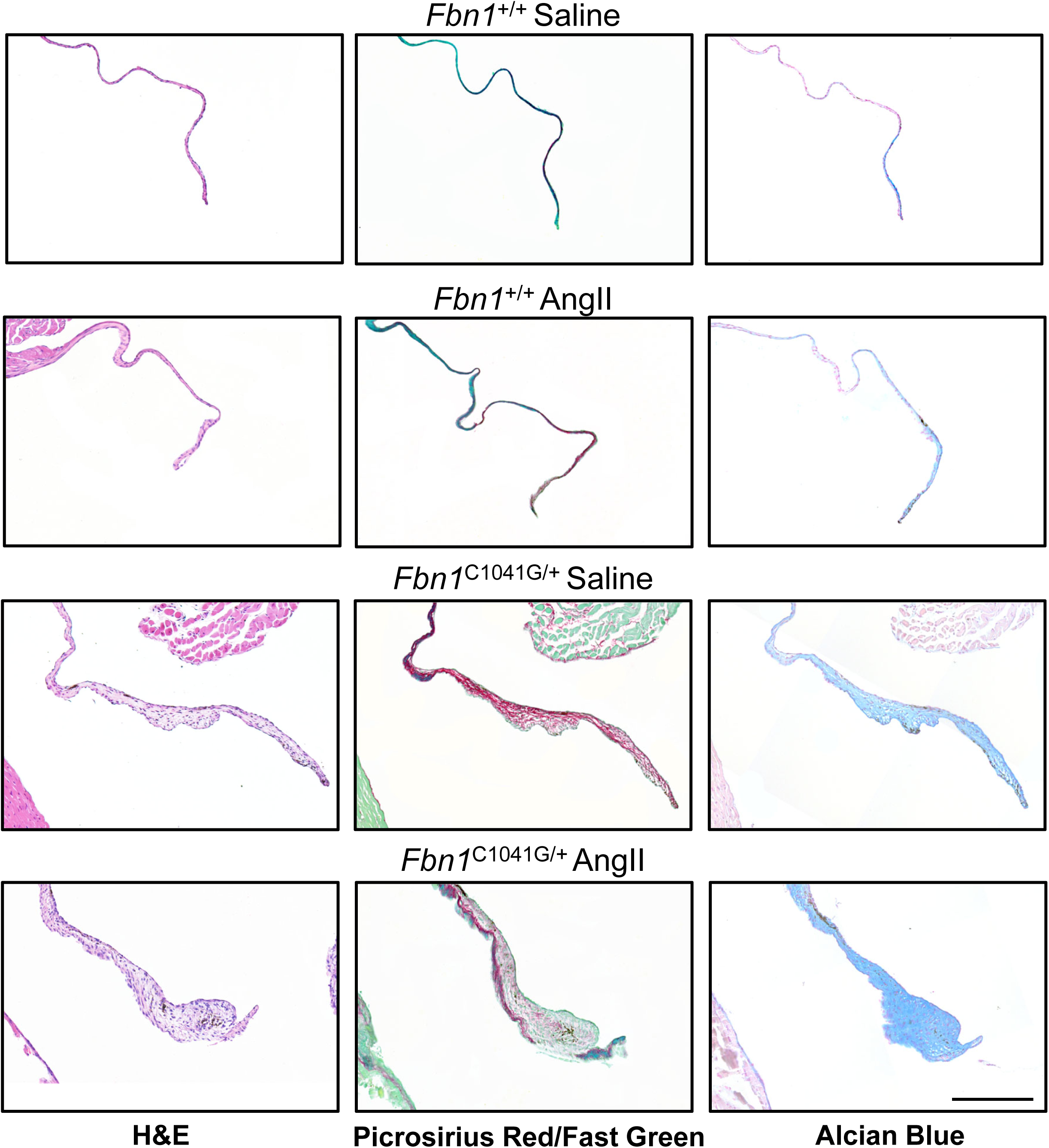
Examples of images of anterior mitral valve leaflet. Examples are for *Fbn1*^+/+^ and *Fbn1*^C1041G/+^ mice infused with either saline or AngII (1,000 ng/kg/min) for 4 weeks. Tissue sections stained with hematoxylin/eosin, picrosirius/red fast green, and alcian blue/nuclear red. Scale bar equals 200 µm.

**Figure S6.**
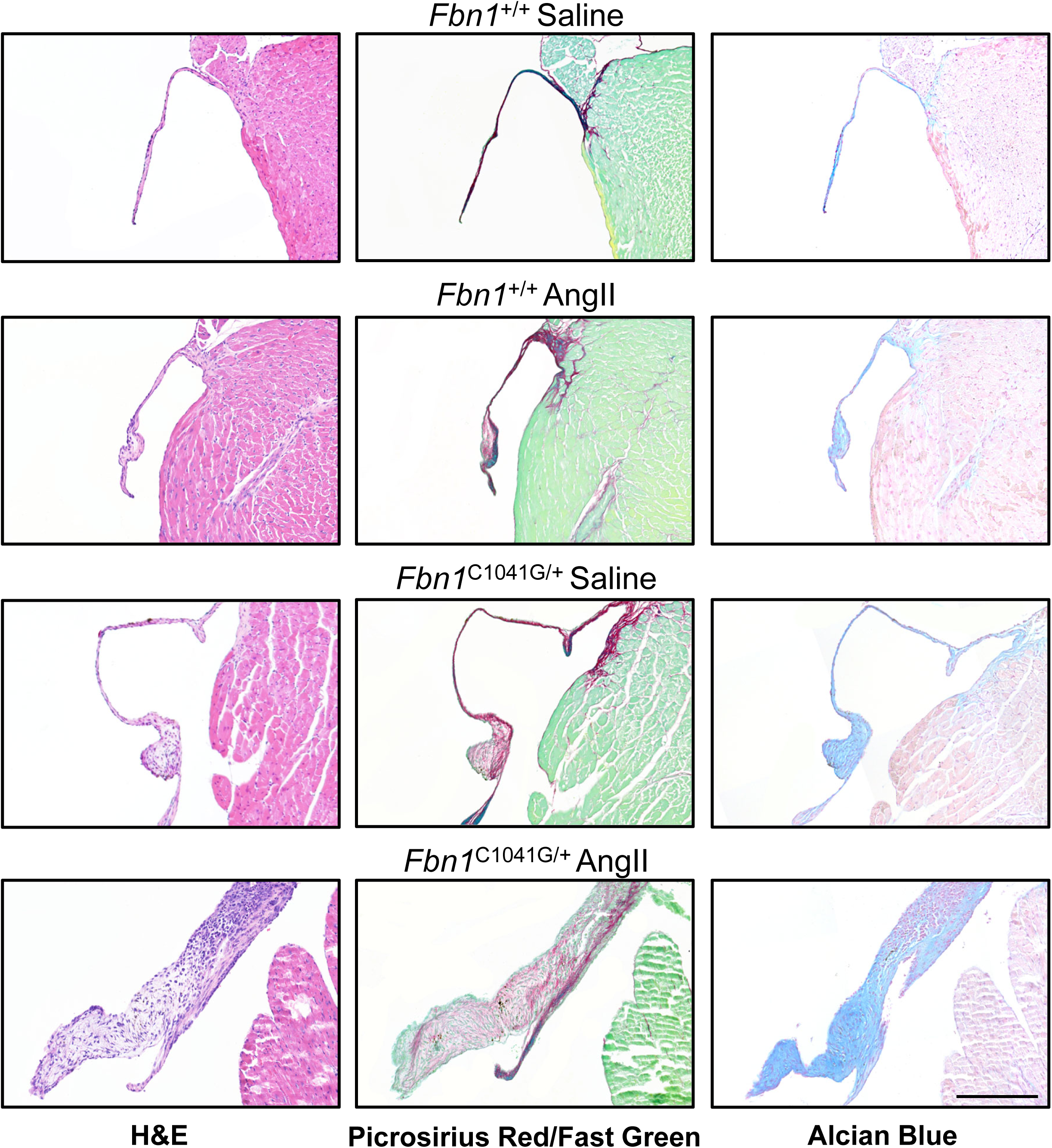
Examples of images of posterior mitral valve leaflet. Examples are for *Fbn1*^+/+^ and *Fbn1*^C1041G/+^ mice infused with either saline or AngII (1,000 ng/kg/min) for 4 weeks. Tissue sections stained with hematoxylin/eosin, picrosirius red/ fast green, and alcian blue/nuclear red. Scale bar equals 200 µm.

**Video S1**. **Representative ultrasound (U/S) videos and images of *Fbn1*^C1041G/+^ with various forms of mitral valve prolapse**. *Fbn1*^+/+^ mouse with normal mitral valve function compared to an *Fbn1*^C1041G/+^ mouse with mitral valve prolapse and regurgitation (yellow arrow) (**A**). *Fbn1*^C1041G/+^ mice with posterior or anterior mitral valve prolapse (**B**). *Fbn1*^C1041G/+^ mice chronically administered AngII with mitral valve prolapse (**C**) and a flailed posterior mitral valve leaflet of an animal that did not survive to the study endpoint (**D**). LA: left atrium, LV: left ventricle, AMVL: anterior mitral valve leaflet, and PMVL: posterior mitral valve leaflet.

**Table S1.**
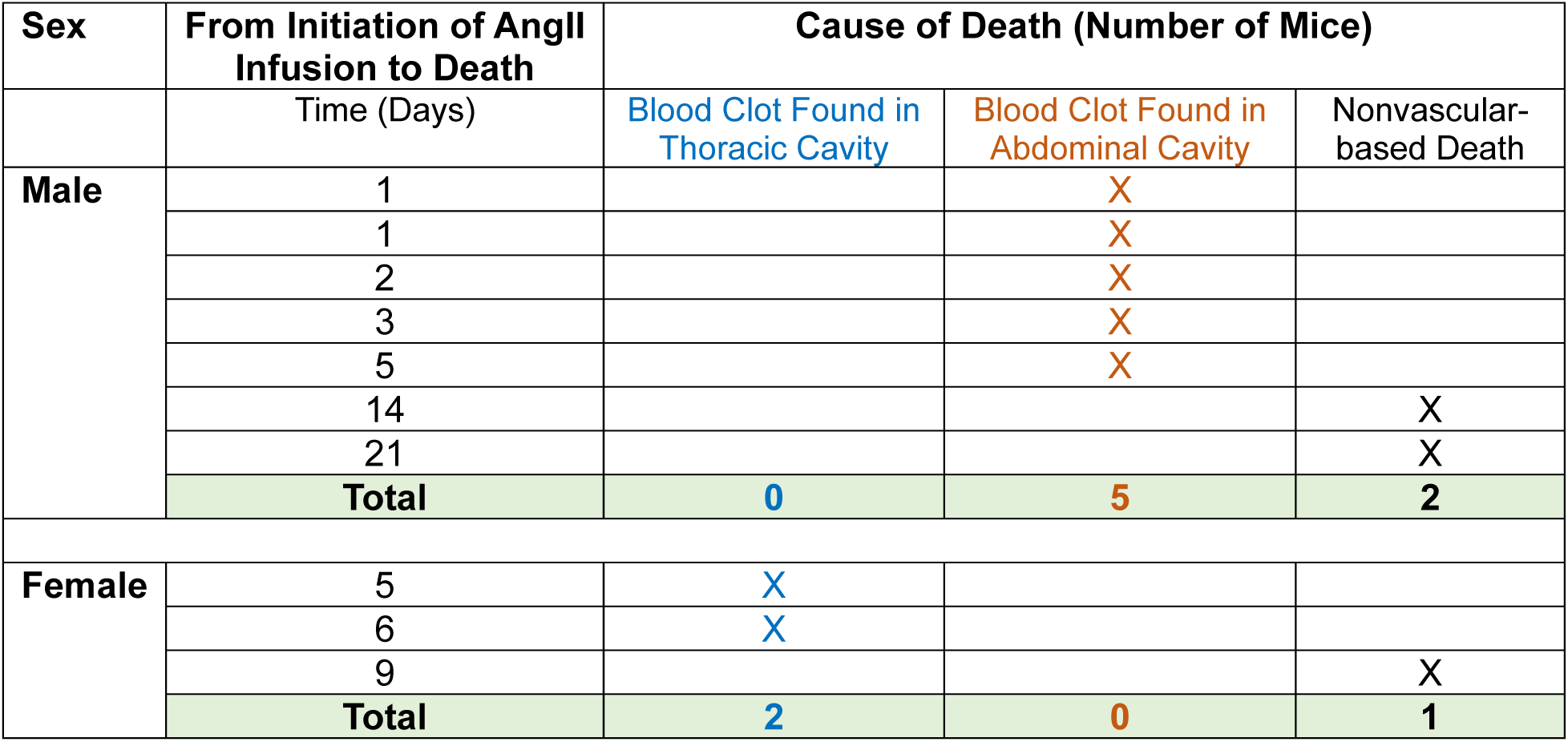
Necropsy on *Fbn1^C1041G/+^* mice represented in Figure 1C-F that died during AngII infusion.

| Sex | From Initiation of AngII Infusion to Death<br>Time (Days) | Cause of Death (Number of Mice) |  |  |
| --- | --- | --- | --- | --- |
|  |  | Blood Clot Found in<br>Thoracic Cavity | Blood Clot Found in<br>Abdominal Cavity | Nonvascular-<br>based Death |
| Male | 1 |  | X |  |
|  | 1 |  | X |  |
|  | 2 |  | X |  |
|  | 3 |  | X |  |
|  | 5 |  | X |  |
|  | 14 |  |  | X |
|  | 21 |  |  | X |
|  | **Total** | **0** | **5** | **2** |
| Female | 5 | X |  |  |
|  | 6 | X |  |  |
|  | 9 |  |  | X |
|  | **Total** | **2** | **0** | **1** |

**Table S2.**
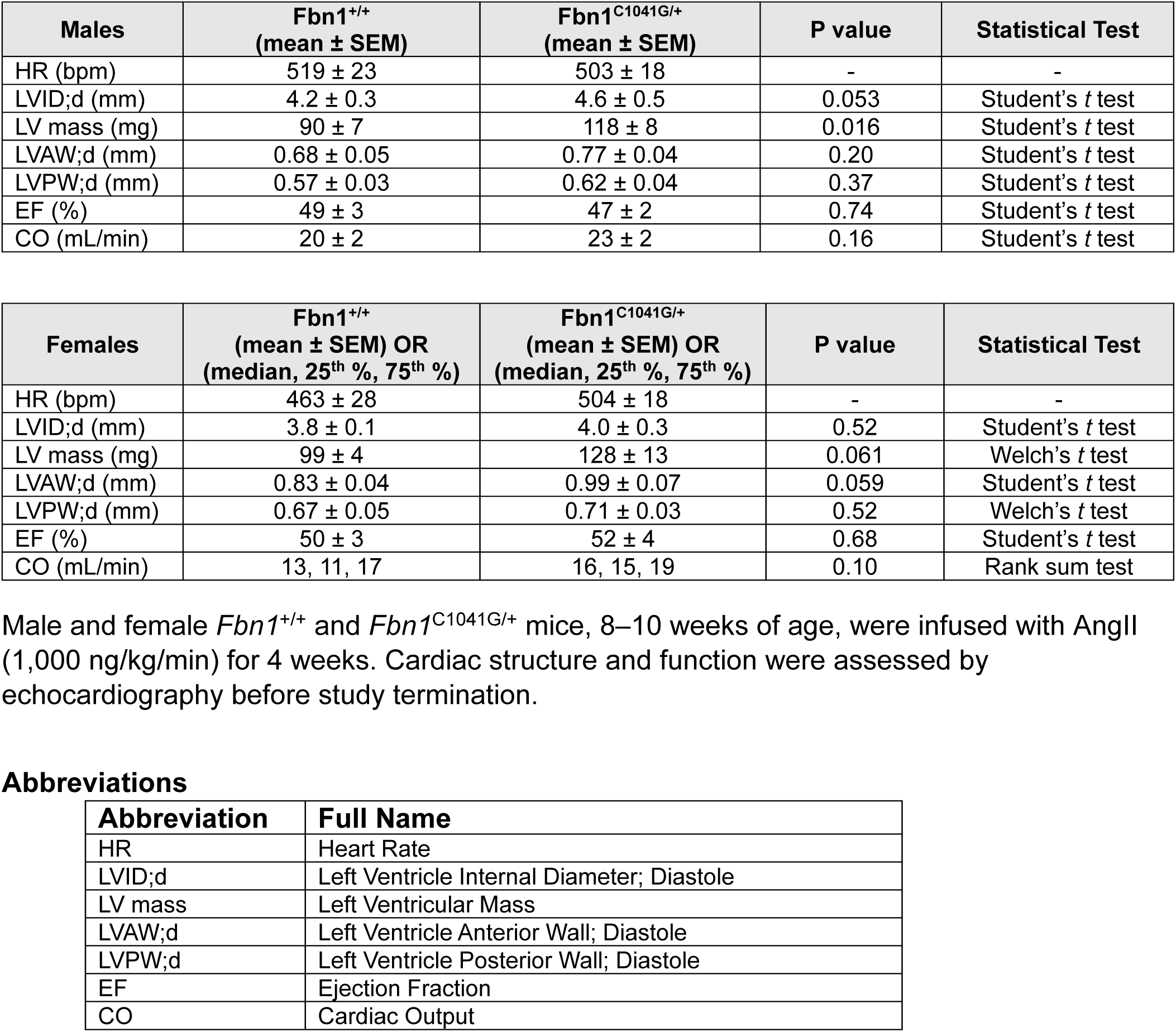
Echocardiography in mice infused with angiotensin II (AngII)

| **Males** | **Fbn1<sup>+/+</sup>**<br>**(mean ± SEM)** | **Fbn1<sup>C1041G/+</sup>**<br>**(mean ± SEM)** | **P value** | **Statistical Test** |
| --- | --- | --- | --- | --- |
| HR (bpm) | 519 ± 23 | 503 ± 18 | - | - |
| LVID;d (mm) | 4.2 ± 0.3 | 4.6 ± 0.5 | 0.053 | Student's *t* test |
| LV mass (mg) | 90 ± 7 | 118 ± 8 | 0.016 | Student's *t* test |
| LVAW;d (mm) | 0.68 ± 0.05 | 0.77 ± 0.04 | 0.20 | Student's *t* test |
| LVPW;d (mm) | 0.57 ± 0.03 | 0.62 ± 0.04 | 0.37 | Student's *t* test |
| EF (%) | 49 ± 3 | 47 ± 2 | 0.74 | Student's *t* test |
| CO (mL/min) | 20 ± 2 | 23 ± 2 | 0.16 | Student's *t* test |

| **Females** | **Fbn1<sup>+/+</sup>**<br>**(mean ± SEM) OR**<br>**(median, 25<sup>th</sup> %, 75<sup>th</sup> %)** | **Fbn1<sup>C1041G/+</sup>**<br>**(mean ± SEM) OR**<br>**(median, 25<sup>th</sup> %, 75<sup>th</sup> %)** | **P value** | **Statistical Test** |
| --- | --- | --- | --- | --- |
| HR (bpm) | 463 ± 28 | 504 ± 18 | - | - |
| LVID;d (mm) | 3.8 ± 0.1 | 4.0 ± 0.3 | 0.52 | Student's *t* test |
| LV mass (mg) | 99 ± 4 | 128 ± 13 | 0.061 | Welch's *t* test |
| LVAW;d (mm) | 0.83 ± 0.04 | 0.99 ± 0.07 | 0.059 | Student's *t* test |
| LVPW;d (mm) | 0.67 ± 0.05 | 0.71 ± 0.03 | 0.52 | Welch's *t* test |
| EF (%) | 50 ± 3 | 52 ± 4 | 0.68 | Student's *t* test |
| CO (mL/min) | 13, 11, 17 | 16, 15, 19 | 0.10 | Rank sum test |

Abbreviations
| **Abbreviation** | **Full Name** |
| --- | --- |
| HR | Heart Rate |
| LVID;d | Left Ventricle Internal Diameter; Diastole |
| LV mass | Left Ventricular Mass |
| LVAW;d | Left Ventricle Anterior Wall; Diastole |
| LVPW;d | Left Ventricle Posterior Wall; Diastole |
| EF | Ejection Fraction |
| CO | Cardiac Output |

### MAJOR RESOURCES TABLES

**Table I.**
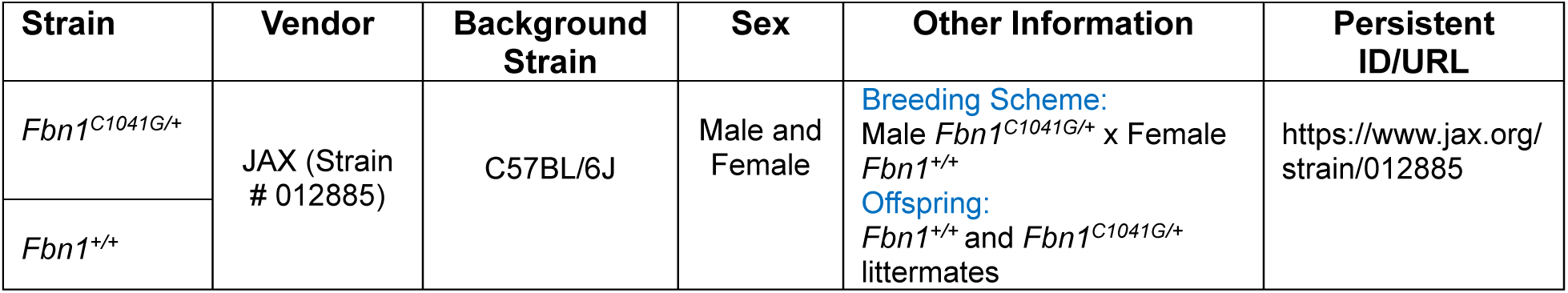
Mouse Models Used in This Study.

**Table II.** Reagents for Histology.

| Reagent | Vendor | Cat # | Working Concentration |
| --- | --- | --- | --- |
| Antigen retrieval solution | Biogenex | HK547-XAK | Ready-to-use |
| Eosin Y | abcam | AB246824 | Ready-to-use |
| Goat serum | Vector Laboratories | MP-7451 | 2.5% vol/vol |
| Hydrogen peroxide | Fisher Scientific | H325-500 | 1% vol/vol |
| Limonene | Medical Chemical Corporation | 6533A | Ready-to-use |
| Mayer's hematoxylin | Electron Microscopy Sciences | 26043-06 | Ready-to-use |
| NovaRED substrate kit | Vector Laboratories | SK-4805 | Ready-to-use |

**Table III.**
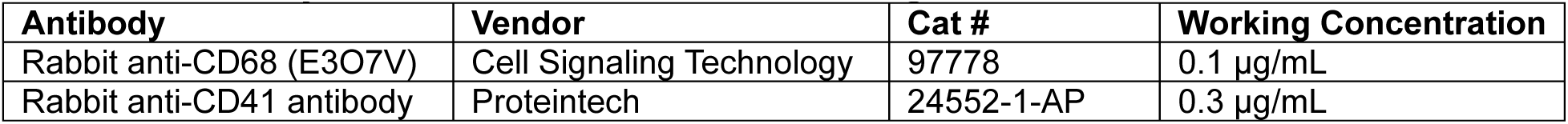
Primary Antibodies for Immunostaining.

**Table IV.**
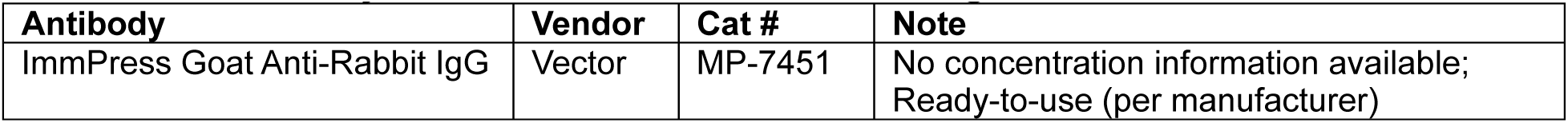
Secondary Antibodies for Immunostaining.

**Table V.** Animal Study Information Following the ARRIVE Essential 10 Figure 1A and S1A.

| Groups | Sex | Age (weeks)* | Number (Start) | Number (Termination) |
| --- | --- | --- | --- | --- |
| *Fbn1*<sup>+/+</sup> PBS | Male | 4-5 | 6 | 6 |
| *Fbn1*<sup>C1041G/+</sup> PBS | Male | 4-5 | 6 | 6 |
| *Fbn1*<sup>C1041G/+</sup> AGT ASO | Male | 4-5 | 6 | 6 |

**Figure 1B and S1B**
| Groups | Sex | Age (weeks)* | Number (Start) | Number (Termination) |
| --- | --- | --- | --- | --- |
| *Fbn1*<sup>+/+</sup> PBS | Female | 4-5 | 6 | 6 |
| *Fbn1*<sup>C1041G/+</sup> PBS | Female | 4-5 | 6 | 6 |
| *Fbn1*<sup>C1041G/+</sup> AGT ASO | Female | 4-5 | 6 | 6 |

**Figure 1C and D**
| Groups | Sex | Age (weeks)* | Number (Start) | Number (Termination) |
| --- | --- | --- | --- | --- |
| *Fbn1*<sup>+/+</sup> Saline | Male | 8-10 | 5 | 5 |
| *Fbn1*<sup>+/+</sup> AngII | Male | 8-10 | 5 | 5 |
| *Fbn1*<sup>C1041G/+</sup> Saline | Male | 8-10 | 6 | 6 |
| *Fbn1*<sup>C1041G/+</sup> AngII | Male | 8-10 | 12 | 5 |

**Figure 1E and F**
| Groups | Sex | Age (weeks)* | Number (Start) | Number (Termination) |
| --- | --- | --- | --- | --- |
| *Fbn1*<sup>+/+</sup> Saline | Female | 8-10 | 6 | 6 |
| *Fbn1*<sup>+/+</sup> AngII | Female | 8-10 | 5 | 5 |
| *Fbn1*<sup>C1041G/+</sup> Saline | Female | 8-10 | 7 | 7 |
| *Fbn1*<sup>C1041G/+</sup> AngII | Female | 8-10 | 10 | 7 |

**Figure 1G**
| Groups | Sex | Age (weeks)* | Number (Start) | Number (Termination) |
| --- | --- | --- | --- | --- |
| *Fbn1*<sup>+/+</sup> Saline | Male | 8-10 | 5 | 5 |
| *Fbn1*<sup>+/+</sup> AngII | Male | 8-10 | 5 | 5 |
| *Fbn1*<sup>C1041G/+</sup> Saline | Male | 8-10 | 5 | 5 |
| *Fbn1*<sup>C1041G/+</sup> AngII | Male | 8-10 | 5 | 5 |

**Figure 1H**
| Groups | Sex | Age (weeks)* | Number (Start) | Number (Termination) |
| --- | --- | --- | --- | --- |
| *Fbn1*<sup>+/+</sup> Saline | Female | 8-10 | 4 | 4 |
| *Fbn1*<sup>+/+</sup> AngII | Female | 8-10 | 6 | 6 |
| *Fbn1*<sup>C1041G/+</sup> Saline | Female | 8-10 | 4 | 4 |
| *Fbn1*<sup>C1041G/+</sup> AngII | Female | 8-10 | 7 | 7 |

**ARRIVE Essential 10 Checklist**
| Item | Application |
| --- | --- |
| Ethics | Approved by the University of Kentucky IACUC (Protocol #: 2018-2967). |
| Sex | Described in each figure. |
| Inclusion criteria | Based on sex, age, body weight, and overall health appearance in each experiment. |
| Exclusion criteria | Based on inconsistent genotypes between pre-study and post-study verification, and medical cases reported by a veterinarian. |
| Sample size | Described in Major Resources Tables |
| Sample size calculation | None |
| Primary endpoint | Mitral valve thickness |
| Randomization | Study mice were numbered and grouped randomly based on their genotype and sex. |
| Blinding | Maximal mitral valve thickness and ultrasound parameters were measured by an investigator blinded to the study group assignment. The data were independently verified by a staff member. |
| Statistical analysis | SigmaPlot 16.0 (SYSTAT Software Inc., CA) |
| Statistical method | Described in each figure legend. |
| Data availability | All numerical data used for figures and supplemental figures are available in the Supplemental Excel File. |

