## Supplementary material for "Angiotensin II Promotes Mitral Valve and Left Atrial Pathology in Fibrillin-1^C1041G/+^ Mice": Video file

### Slide 1
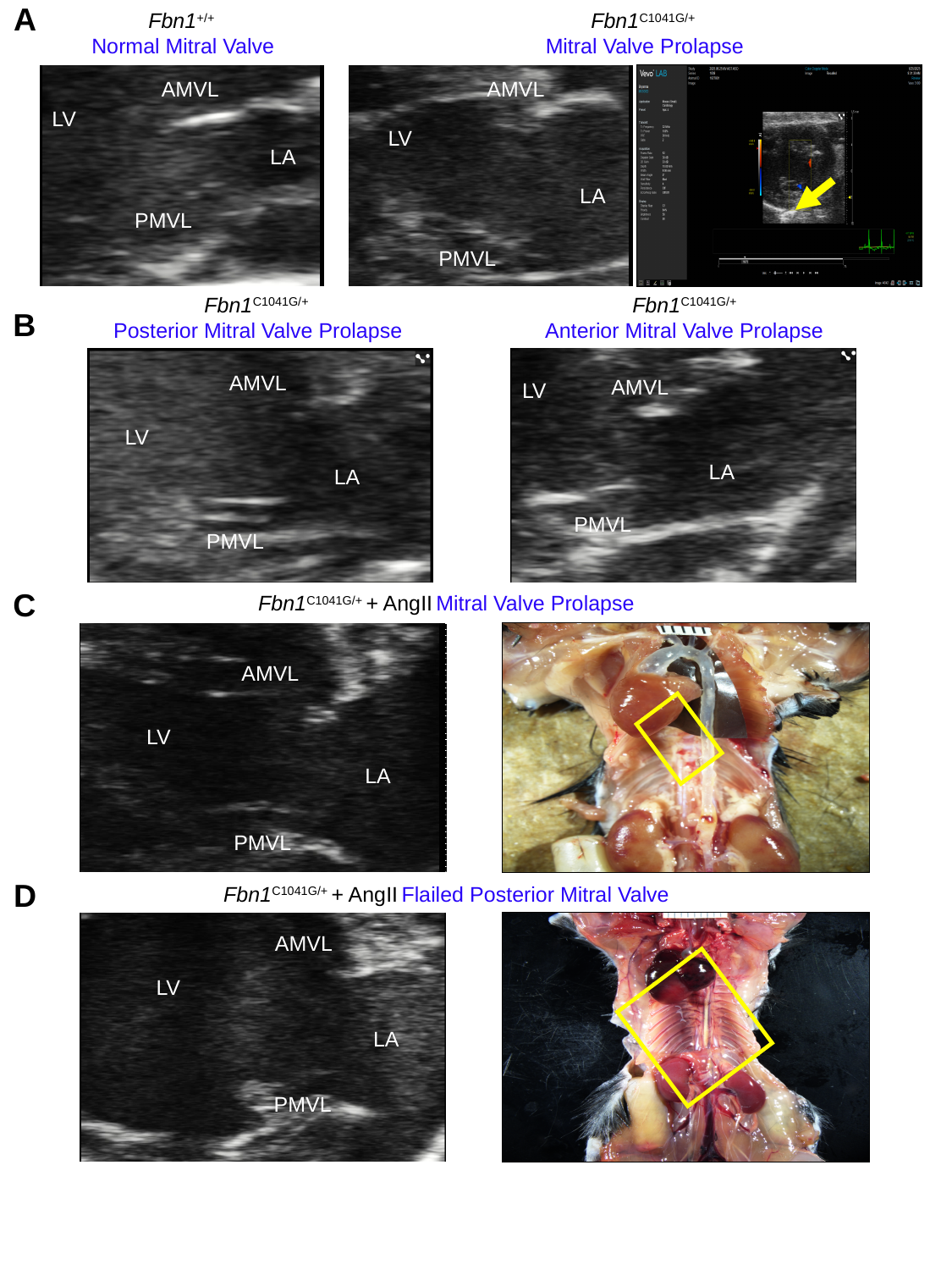

A
Fbn1^C1041G/+^
Mitral Valve Prolapse
Fbn1^+/+^
Normal Mitral Valve
AMVL
AMVL
LV
LV
LA
LA
PMVL
PMVL
Fbn1^C1041G/+^
Anterior Mitral Valve Prolapse
Fbn1^C1041G/+^
Posterior Mitral Valve Prolapse
B
AMVL
AMVL
LV
LV
LA
LA
PMVL
PMVL
C
Fbn1^C1041G/+^ + AngII Mitral Valve Prolapse
AMVL
LV
LA
PMVL
D
Fbn1^C1041G/+^ + AngII Flailed Posterior Mitral Valve
AMVL
LV
LA
PMVL
